# Improved Genome Editing by an Engineered CRISPR-Cas12a

**DOI:** 10.1101/2022.09.02.506401

**Authors:** Enbo Ma, Kai Chen, Honglue Shi, Elizabeth C. Stahl, Ben Adler, Junjie Liu, Kaihong Zhou, Jinjuan Ye, Jennifer A. Doudna

## Abstract

CRISPR-Cas12a is an RNA-guided, programmable genome editing enzyme found within bacterial adaptive immune pathways. Unlike CRISPR-Cas9, Cas12a uses only a single catalytic site to both cleave target double-stranded DNA (dsDNA) (*cis*-activity) and indiscriminately degrade single-stranded DNA (ssDNA) (*trans*-activity). To investigate how the relative potency of *cis*- versus *trans*-DNase activity affects Cas12a-mediated genome editing, we first used structure-guided engineering to generate variants of *Lachnospiraceae bacterium* Cas12a (LbCas12a) that selectively disrupt *trans*-activity. The resulting engineered mutant with the biggest differential between *cis*- and *trans*-DNase activity *in vitro* showed minimal genome editing activity in human cells, motivating a second set of experiments using directed evolution to generate additional mutants with robust genome editing activity. Notably, these engineered and evolved mutants had enhanced ability to induce homology-directed repair (HDR) editing by 2-18-fold depending on the genomic locus. Finally, we found that a site-specific reversion mutation produced improved Cas12a (iCas12a) variants with superior genome editing efficiency at genomic sites that are difficult to edit using wild-type Cas12a. This strategy of coupled rational engineering and directed evolution establishes a pipeline for creating improved genome editing tools by combining structural insights with randomization and selection. The availability of experimental and predicted structures of other CRISPR-Cas enzymes will enable this strategy to be applied to improve the efficacy of other genome editing proteins.

## INTRODUCTION

CRISPR (clustered regularly interspaced short palindromic repeats)-Cas (CRISPR-associated proteins) systems provide versatile adaptive immunity in prokaryotes as well as programmable genome editing tools in eukaryotes (1–4). Both Cas9 and Cas12a proteins have been widely deployed for programmable genome editing, each with distinct properties that could impact editing outcomes. A potentially important difference in the biochemical behavior between Cas9 and Cas12a is the ability of Cas12a to carry out dual nuclease activities: cleaving target double-stranded DNA (dsDNA) guided by CRISPR RNA (crRNA) (*cis*-activity) and non-specifically degrading single-stranded DNA (ssDNA) following crRNA-guided target DNA binding and cutting (*trans*-activity) (Figure 1A). This *trans*-ssDNase activity has been harnessed for Cas12a-mediated nucleic acid detection and viral diagnosis (5–8). However, how and whether the *trans*-nuclease activity could impact genome editing outcomes are not yet fully understood.

**Figure 1.**
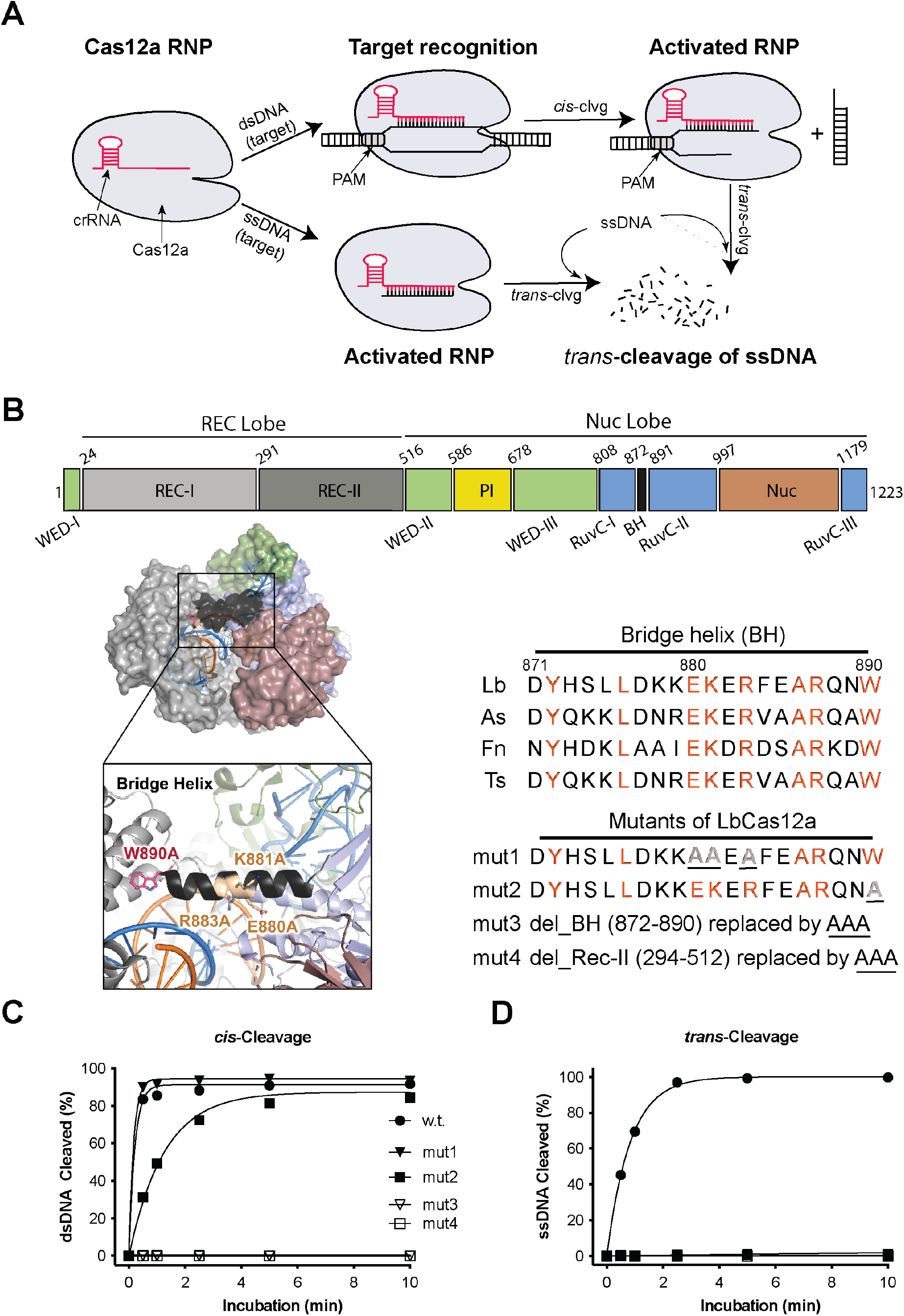
Importance of the bridge helix (BH) of LbCas12a protein in regulating its nuclease activities. **A**. Illustration of *cis*- and *trans*-cleavage activities of Cas12a proteins. *Trans*-activity of Cas12a RNP can be activated by either direct binding to target ssDNA or after processing its target dsDNA. **B**. Schematic presentation of LbCas12a protein. Domain assignment for LbCas12a (upper panel), protein structure (PDB ID: 5XUS) highlighting the bridge helix in LbCas12a (lower panel, left), and bridge helix sequences of different Cas12a orthologs and designed mutations from LbCas12a (lower panel, right). Point mutations are underlined, and deletions are replaced with a triple alanine sequence (AAA) which is also underlined. **C** and **D**. *In vitro* kinetic studies of *cis*- and *trans*-cleavage activities by wild-type (WT) and four designed mutants, mut1-4. Each data point is the average of two to three parallel experiments.

Cas12a comprises three structural regions: the recognition (REC) lobe, the nuclease (Nuc) lobe (including the RuvC, bridge helix (BH), and Nuc domains), and a connection region (including the Wedge (WED), and PAM-interacting (PI) domains) between the two lobes (9, 9–11) (Figure 1A). It has been shown that Cas12a uses a single active site located in the RuvC domain to cleave both strands of the target dsDNA in *cis*-activity as well as non-specific ssDNA in *trans*-activity, following a well-defined order in the reaction steps (11–14). When CRISPR-Cas12a binds the target DNA, the spacer region of the crRNA forms Watson-Crick base pairs with its complementary target strand (TS) in the target DNA, leaving the non-target strand (NTS) unpaired, to form an R-loop. Upon R-loop formation, the non-target strand (NTS) is immediately accessed by the active site and cleaved or nicked, and further trimmed by five nucleotides (nts) to make the gap big enough for the TS to reach the active site and be cut (13, 14). After cleavage, the TS is released from the catalytic site to enable *trans*-cleavage of non-specific ssDNA. It has been shown that *trans* ssDNA cleavage requires NTS cleavage and can be accelerated by both NTS trimming and TS cleavage (13, 14). Given this sequential pathway of Cas12a reaction steps, it seemed possible that Cas12a’s *trans*-cutting activity might be inhibited or reduced with minimal effect on target-specific *cis*-cleavage activity.

To investigate the effect of disruptions in Cas12a’s dual nuclease activity on genome editing, we employed a combination of structure-guided engineering and directed evolution to identify Cas12a variants that minimize *trans*-DNase activity while retaining robust *cis*-dsDNA cutting behavior. In principle, Cas12a proteins with reduced *trans*-activity might improve DNA integration efficiency as an editing outcome following homology-directed repair (HDR) since *trans*-activity could potentially degrade ssDNA donor templates used for HDR.

A previous biochemical and single-molecule study showed that perturbations at the bridge helix (BH), which resides between the RuvC and REC-II domains and is responsible for RuvC activation, minimally impacted Cas12a’s *cis*-cleavage activity but reduced the NTS trimming activity in *Franicisella novicida* Cas12a (FnCas12a) (15). We therefore hypothesized that mutations in the BH region might disfavor *trans*-cutting while maintaining target-specific *cis*-cleavage activity. Using structural-based engineering, we found that a W890A mutation in the BH of *Lachnospiraceae bacterium ND2006* Cas12a (LbCas12a) abolished *trans*-activity while modestly reducing *cis*-activity. Combined with beneficial mutations identified by directed evolution, we produced Cas12a variants with both high-level *cis*-nuclease activity and minimized *trans*-ssDNase activity. These new variants robustly induce genome edits in human cells, with elevated homology-directed repair (HDR) efficiency. Surprisingly, the restoration of W890 in these beneficial variants created improved Cas12a (iCas12a) proteins that possess significantly enhanced *cis*- and *trans*-activity *in vitro* and improved genome editing activity in human cells relative to the original Cas12a protein. This study demonstrates that the dual activities of CRISPR-Cas12a can be altered to enhance its application for genome manipulation.

## MATERIALS AND METHODS

### Preparation of designed CRISPR-Cas12a mutants

The mutants of CRISPR *Lachnospiraceae bacterium* Cas12a (LbCas12a) bearing mutations in the bridge helix (BH) domain were generated by site-directed mutagenesis polymerase chain reaction (PCR) in the presence of a given wild-type LbCasd12a plasmid, two corresponding primers (synthesized by IDT) and KOD polymerase (Sigma-Aldrich). Specifically, the reaction was carried out in 25μl reaction containing 10ng of wild-type LbCas12a plasmid and 0.75μl of 10μM primers containing desired mutations. After PCR, the reaction was treated with 1μl of DpnI (BioLabs, New England) for 1 hour at 37°C before transformation. The wild-type LbCas12a plasmid is a home-made pET-based expression vector containing an N-terminal His_10_-tag, maltose-binding protein (MBP) and TEV protease cleavage site (QB3 MacroLab, UC Berkeley). The sequences of all of the plasmid constructs are confirmed via Sanger sequencing (UC Berkeley DNA Sequencing Facility).

### Generation of new LbCas12a mutants via directed evolution

To engineer the mutant LbCas12a, mut2, for higher nuclease activities, we performed directed evolution using a bacteria selection system (Figure 3A). First, we made a Chloramphenicol-resistant (CAM^+^) bacterial expression construct containing both point mutation of W890A (LbCas12a-mut2) and a guide crRNA that targets its complementary sequence containing protospacer adjacent motif (PAM) of TTTG in the *ccdB* toxin gene of the selection plasmid. This inducible *ccdB* gene is constructed in a plasmid with Ampicillin-resistant (Amp^+^) gene and its expression is controlled by the arabinose-promoter pBAD for a positive selection. To simplify the engineering, we divided the LbCas12-mut2 protein into 3 fragments (Figure 3B): fragment 1 (R1) covers1-532; fragment2, 507-820 (R2); and fragment3, 797-1228 (R3). To generate mutant libraries, we first performed error-prone PCR on each target fragment with an error rate of 6- to 9-nucleotide mutations per kilobase and the error-prone PCR products are then used to replace the corresponding parent fragment in the founder plasmid of LbCas12a-mut2. The error-prone PCR was carried out with the ThermoTaq DNA polymerase (M0267S, NEB) in a reaction of 100μl with 10μl of 10X ThermoPol reaction buffer, 2μl of 10mM primers, 2.4μl of 10mM MnCl_2_, 32ng of template plasmid and 1μl of ThermoTaq DNA Polymerase. Two nanogram of the error-prone library plasmid DNA were electroporated in 50μl of competent cells made from *E. coli* strain *BW25141(DE3)* that contains the selection plasmid encoding the arabinose-inducible *ccdB* toxin gene.

After recovery of the electroporated bacteria in 2ml of SOB for 50 min at 37°C, 5 μl of the bacteria culture was plated onto a Petri agar-dish containing only CAM (as control) and the remainder culture was plated on another Petri agar-dish containing both arabinose and CAM. Positive colonies that grew on the plates containing both arabinose and CAM were collected and replated. Plasmids of individual colonies from the replated plate were then prepared and sequenced. In this study, two rounds of section were carried out.

### Protein expression and purification

The protocol for purification of LbCas12a proteins has been described elsewhere (5). Simply, all of Cas12a proteins were expressed in *E. coli* Rosetta (DE3) cells (Sigma-Aldrich) cultured in Terrific Broth (TB) medium (Thermo Fisher Scientific) supplemented with the antibiotics of ampicillin and chloramphenicol. The culture was carried out at 37°C after inoculation with overnight-cultured starter at a 1:40 ratio. When optical density (OD_600_) of the culture reached 0.6-0.8, the bacteria was induced by the addition of isopropyl β-D-1-thioglalacctopyranoside (IPTG) to a final concentration of 0.1 mM and incubated overnight at 16°C. To purify the Cas12a proteins, the cultured cells were harvested and resuspended in Lysis Buffer (LB: 50 mM Tris-HCl, pH 7.5, 500 mM NaCl, 5% (v/v) glycerol, 1 mM TCEP, 0.5 mM PMSF and 0.25 mg/ml lysozyme as well as a cOmplete™ Protease Inhibitor Cocktail Tablet (Millipore Sigma) for every 50 ml), disrupted by sonication and centrifuged for 60 min at x18,000 rcf. The supernatant was incubated with Ni-NTA resin for 60 minutes to pull down the His-tagged target protein. After the cleavage with TEV protease (home-made) overnight at 4°C, the target proteins were separated from His-tagged MBP via HiTrap Ni-NTA column (GE) and further purified over a HiTrap Heparin HP column (GE) for cation exchange chromatography. The final gel filtration step (Superdex 200, GE) was carried out in gel-filtration buffer containing 20 mM Tris-HCl, pH 7.5, 200 mM NaCl, 5% (v/v) glycerol and 1 mM TCEP.

### Nucleic acid preparation

All of the DNA and RNA oligos used in this study were purchased from Integrated DNA Technologies, Inc. (IDT) and HPLC or PAGE-purified. crRNAs purchased from IDT possess chemical modifications at 3’- or 5’-ends, according to the IDT guidance, to improve their stability and editing efficiency in cells.

### DNA cleavage assays

Typical *cis*-cleavage assays (otherwise will be stated) were carried out with 30nM protein, 36nM of crRNA and 5-10nM of 5’-FAM-labeled target DNA in the cleavage buffer consisting of 20mM HEPES (pH 7.5), 150mM KCl, 10mM MgCl_2_, 1% glycerol and 0.5mM TCEP. Specifically, the protein and a guide crRNA were first incubated for 15 min at room temperature to form RNPs and then incubated at 37°C for certain duration of time after addition of labeled target DNA. For *transcleavage* assays, the reaction conditions were the same as *cis*-cleavage reactions except that 45nM unlabeled target dsDNA instead of labeled target dsDNA were used. After incubation for 30 min at 37°C, a labeled random ssDNA was added to the reactions and the incubation continued for certain duration of time. The reactions were quenched with DNA loading buffer (45% formamide and 15 mM EDTA, with trace amount of xylene cyanol and bromophenol blue). After denatured at 95°C for 3 min, the cleavage products were separated in 15% urea-PAGE gel and quantified with Typhoon (Amersham, GE Healthcare).

For radiolabeled cleavage assays, the substrates used are 5’-end-labeled with T4 PNK (NEB) in the presence of gamma ^32^P-ATP. For dsDNA substrates, the target or non-target strand is first 5’-end-labeled and then annealed with excess (1.2 times) corresponding strand. The concentrations of Cas12a, guide RNA and ^32^P-labeled substrates used in the reaction are 30nM, 36nM and 2-4nM (unless otherwise stated), respectively. Reactions were incubated for certain minutes at 37°C and quenched with DNA loading buffer for 2-3 min at 90°C. The substrates and products were resolved by 15% urea-denaturing PAGE gel and quantified with Typhoon (Amersham, GE Healthcare).

For time course studies, fitting curve of a cleavage assay was generated by the formula of Y = Y_max_ × (1 – e^−kt^), where Y_max_ is the pre-exponential factor, k is the rate constant (min^−1^), and t is the reaction time (min).

### Cell culture and genomic editing

To evaluate the genome editing activity of different LbCas12a mutant proteins, we first assembled their RNPs with corresponding crRNAs before performing nucleofection with HEK293T cells or tdTomato neural progenitor cells (NPCs). Specifically, the protein (100 pmol) and the crRNA (120 pmol) with a mole ratio of 1:1.2 were first incubated for 15-25 min at room temperature to form RNPs, and then mixed with a ssDNA electroporation enhancer (80 pmol, purchased from IDT) with a mole ratio of 1:0.8 (RNP:enhancer) (the electroporation enhancer was used for all the cell editing experiments based on nucleofection unless otherwise noted). For HDR experiments, ssDNA donor template (100 pmol) was further added to the RNP solution with a mole ratio of 1:1 (RNP:donor ssDNA). Lonza SF (for HEK293T cells) and P3 (for tdTomato NPCs) buffers were used for the preparation of nucleofection mixtures.

HEK293T cells (UC Berkeley Cell Culture Facility) were cultured using Dulbecco’s Modification of Eagle’s Medium (DMEM) with L-glutamine, 4.5g/L glucose and sodium pyruvate (Corning) plus 10% FBS, and penicillin and streptomycin (Gibco). Nucleofection of HEK293T cells with RNPs was performed using Lonza (Allendale, NJ) SF cell kits in an Amaxa 96-well Shuttle system with a program code CM-130. Each nucleofection reaction consisted of approximately 2.0 × 10^5^ cells and 100 pmol RNP with a total volume of 25μl in the supplemented SF buffer according to the Lonza protocol. After nucleofection, 75μl of growth media was added to each nucleofection cuvette to transfer the cells to 12-well tissue culture plates with a total culture volume of 1ml/well. For HDR experiments, HDR enhancer (Alt-R™ HDR Enhancer V2, 0.69 mM stock in DMSO, purchased from IDT) with a final concentration of 0.33uM was added to the culture media. After incubation at 37°C for 24 hours, the cell culture media was refreshed. The HEK293T cells were harvested for analysis after further incubation at 37°C for 72 hours.

NPCs were isolated from embryonic day 13.5 Ai9-tdTomato homozygous mouse brains(16). Cells were cultured as neurospheres in NPC medium: DMEM/F12 with glutamine, Na-Pyruvate, 10 mM HEPES, non-essential amino acid, penicillin and streptomycin (100x), 2-mercaptoethanol (1,000x), B-27 without vitamin A, N2 supplement, and growth factors, bFGF and EGF (both 20 ng/ml as final concentration). NPCs were passaged using MACS Neural Dissociation Kit (Papain, CAT# 130-092-628) following manufacturer’s protocol. bFGF and EGF were refreshed every three days and cells were passaged every six days. The NPC line was authenticated by immunocytochemistry marker staining for Nestin and GFAP. Nucleofection of NPCs with RNP was performed using Lonza (Allendale, NJ) P3 cell kits in an Amaxa 96-well Shuttle system with a program code EH-100. Each nucleofection reaction consisted of approximately 2.5 × 10^5^ cells and 100 pmol RNP with a total volume of 20μl in the supplemented P3 buffer according to the Lonza protocol. After nucleofection, 80μl of growth media was added to the nucleofection cuvette and 15μl of NPC culture was then transferred to 96-well tissue culture plates pre-coated (using laminin, fibronectin, and poly-DL-ornithine), with a total culture volume of 100μl/well. After incubation at 37°C for 72 hours, the cell culture media was refreshed. The NPCs were harvested for analysis after further incubation at 37°C for 72 hours.

For direct RNP delivery experiments with NPCs, RNPs were pre-assembled by mixing LbCas12a mutant proteins (100pmol) and the corresponding crRNA (120pmol) in a phosphate buffer (25 mM NaPi, 300 mM NaCl, 200 mM trehalose, pH7.5) with a total volume of 15μl at room temperature, and then added to NPC culture in 96-well plates 36 hours after cell seeding (starting with 2 × 10^4^ cells/well). After incubation at 37°C for 36 hours, the cell culture media was refreshed. The NPCs were harvested for analysis after further incubation at 37°C for 96 hours.

For genomic DNA extraction, the media was removed by aspiration and 100μl of Quick Extraction solution (Epicentre, Madison, WI) was added to lyse the cells (65°C for 20 min and then 95°C for 20 min). The concentration of genomic DNA was determined by NanoDrop and the cell lysate was stored at −20°C. tdTomato-positive NPCs were analyzed by flow cytometry.

To fully investigate target editing (TE) and HDR of each LbCas12a protein on a genomic DNA target in HEK293T cells, target amplicons were PCR-amplified in the presence of corresponding primers which were designed to have no overlap with their corresponding donor ssDNA sequence in the case of HDR. The PCR products were purified with magnetic-beads (Berkeley Sequencing Core Facility) before being subjected to next generation sequencing (NGS) with MiSeq (Illumina) at 2×300bp with a depth of at least 10,000 reads per sample.

The paired-end Illumina sequencing reads were trimmed using the BBDuk tool in Geneious Prime (https://www.geneious.com/prime) with a minimum quality of 20 and a minimum length of 20, and then merged using the BBmerge tool in Geneious Prime. The merged reads were then subjected to CRISPResso2 (https://github.com/pinellolab/CRISPResso2) to quantify the rate of indels and HDR with the two following commands, respectively:

*CRISPResso --fastq_r1 MERGED_READS --amplicon_seq AMPLICON_SEQUENCE -- guide_seq GUIDE_SEQUENCE -n nhej -wc -5 -w 9 --plot_window_size 20 -o OUTPUT_FILE*
*CRISPResso --fastq_r1 MERGED_READS --amplicon_seq AMPLICON_SEQUENCE -- guide_seq GUIDE_SEQUENCE -e DONOR_SEQUENCE -wc -5 -w 9 --plot_window_size 20 -o OUTPUT_FILE*

where *--fastq_r1* is followed by the path of the input merged fastq file, *--amplicon_seq* is followed by the full sequence of amplicon, *--guide_seq* is followed by the spacer sequence of the guide RNA, *--e* is followed by the expected amplicon sequence after successful HDR, *-wc* is followed by the quantification window center relative to the 3’ end of the spacer sequence, *-w* is followed by the size of the quantification window and *--plot_window_size* is followed by the size of the quantification window for visualizing the indels. The position and size of the quantification window were adjusted by the cleavage patterns of Cas12a mutants in this study.

## RESULTS

### *Cas12a bridge helix mutations differentially affect cis*- versus *trans-DNA cleavage*

Sequence alignment of the bridge helix (BH) segments from several Cas12a orthologs revealed a high sequence identity (>40%) with several fully conserved residues (Figure 1B). For instance, residues D877/E880 and R883/R887 in the BH of Cas12a from *Lachnospiraceae bacterium ND2006* (LbCas12a) build an acid-base network with residues K940 and E939 in the RuvC domain, respectively; and another residue, W890, residing at the end of BH, is anchored in a hydrophobic cavity of the REC-II domain and involved in regulating the open-closed conformational change of the whole protein (10, 15, 17). Data from mutagenesis and biochemical assays have shown that the disruption of the BH helical structure or the prevention of anchoring BH to the REC-II domain in FnCas12a minimally reduces the *cis*-cleavage activity but impairs the NTS trimming activity (15). Furthermore, the NTS trimming for gap formation was shown to be a critical conformational prerequisite for the subsequent *trans*-cleavage activity (13, 14). Therefore, we rationalized that mutations in the BH might help us to dissect the *cis*- and *trans*-activities of Cas12a proteins.

We designed 3 mutations in the BH domain of LbCas12a. Two of them are the point-mutations at these highly conserved amino acid positions, named mut1 (E880A, K881A, and R883A) and mut2 (W890A), and another mutant with the entire BH replaced by a triple alanine sequence (AAA), named mut3 (ΔY872-W890). As the REC-II domain is known to participate in the dynamic change and to directly interact with BH domain during the activity of Cas12a (11, 17), an additional mutant, mut4 (ΔY294-T512), with the Rec-II domain replaced by AAA, was also included in our original test (Figure 1B).

Proteins from the four designed mutants were purified (Supplementary Figure S1A) and used to perform *in vitro* cleavage assays using a 45nt CRISPR RNA (crRNA) containing a 21-base direct repeat region (loop domain) and a 24-base protospacer region complementary to the center of a 55-base pair (bp) dsDNA substrate in which the target region flanks the protospacer adjacent motif (PAM) of TTTA (Supplementary Figure S1B). We also tested target-activated ssDNA cleavage by these mutants and compared them to the original wildtype enzyme. The biochemical assays showed that although bridge-helix mutants mut1 and mut2 retained robust *cis*-cleavage activity on dsDNA substrate, their *trans*-ssDNase activity was substantially reduced or diminished (Figure 1C,D, Supplementary Figure S1C-F). However, mut3 and mut4, containing either BH or REC-II domain truncations, showed no detectable *cis*- or *trans*-cleavage activity for either dsDNA or ssDNA substrates (Figure 1C,D, Supplementary Figure S1C-F), indicating that the domain deletions completely disrupt protein conformations required for the catalytic activities, resulting in the loss of function.

As Cas12a utilizes a single RuvC domain to cleave sequentially the two strands of dsDNA and then catalyze *trans*-ssDNA cutting (5, 14, 18, 19), we tried to understand how the dual activities of the RuvC domain are regulated by kinetic analyses of LbCas12a proteins with mutations in its bridge helix domain. From four variants that were tested (mut1-4), we chose mut2 LbCas12a (a point mutation of W890A) for further study due to its relatively robust *cis*-activity and minimal *trans*-activity (Figure 1C, Supplementary Figure S1C-F). Kinetic analysis of *cis*-cleavage was conducted using fluorescently 5’-end-labeled target (TS) or non-target strand (NTS) dsDNA substrates. Although the mutation of W890A in mut2 resulted in a ~50% reduction in initial rate (3.2 min^−1^ vs. 7.1 min^−1^) of non-target strand (NTS) cutting relative to the wildtype (WT) LbCas12a, the mut2-catalyzed TS cutting rate was 10% that of the WT LbCas12a (0.5 min^−1^ vs. 5.7 min^−1^) (Figure 2A, Supplementary Figure S2A,B). This effect was even more evident with ssDNA substrates (Figure 2B, Supplementary Figure S2C,D). Analysis of the NTS and TS cleavage products showed that the mut2-generated NTS cleavage site was shifted by 2-3 nucleotides away from the PAM (Figure 2C) with reduced NTS trimming (Supplementary Figure S2A), relative to the wildtype protein NTS cleavage product, while the cleavage site on TS remained identical between these two proteins (Figure 2C). These results are consistent with previous biochemical studies of BH mutations in FnCas12a (15, 19). These findings support a ‘substrate-occlude’ in which the altered NTS cleavage positions and reduced NTS trimming interfere with the subsequent entry of TS and non-specific ssDNA to the catalytic site, resulting in reduced TS *cis*-cleavage and diminished *trans*-activity (14).

**Figure 2.**
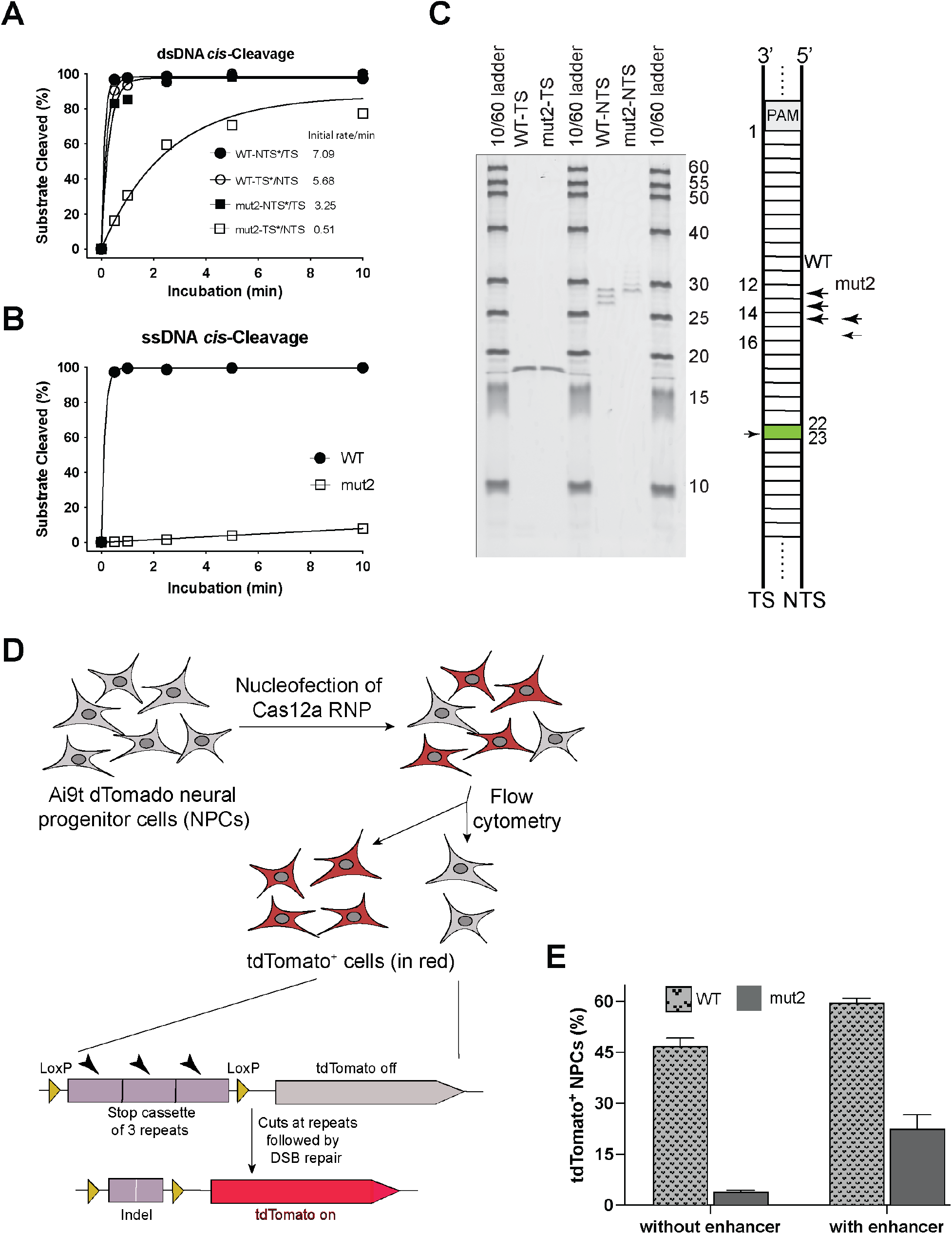
Mutational effect of W890A on the nuclease activities of LbCas12a. **A** and **B**. Kinetic studies of the *cis*-cleavage activities on dsDNA (A) and ssDNA (B) by WT and mut2 (W890A). TS=target strand; NTS= nontarget strand; *=labelled strand. **C**. Mutational effect of W890A on the cleavage sites of NTS of dsDNA. **D**. Genome editing workflow using tdTomato neural progenitor cells (NPCs) from Ai9 mice. The tdTomato gene will be turned on when editing happens to remove the stop cassette. **E**. Genome editing of NPCs by WT and mut2. Editing level is reflected by the percentage of tdTomato-positive NPCs (n = 3, means ± SD).

**Figure 3.**
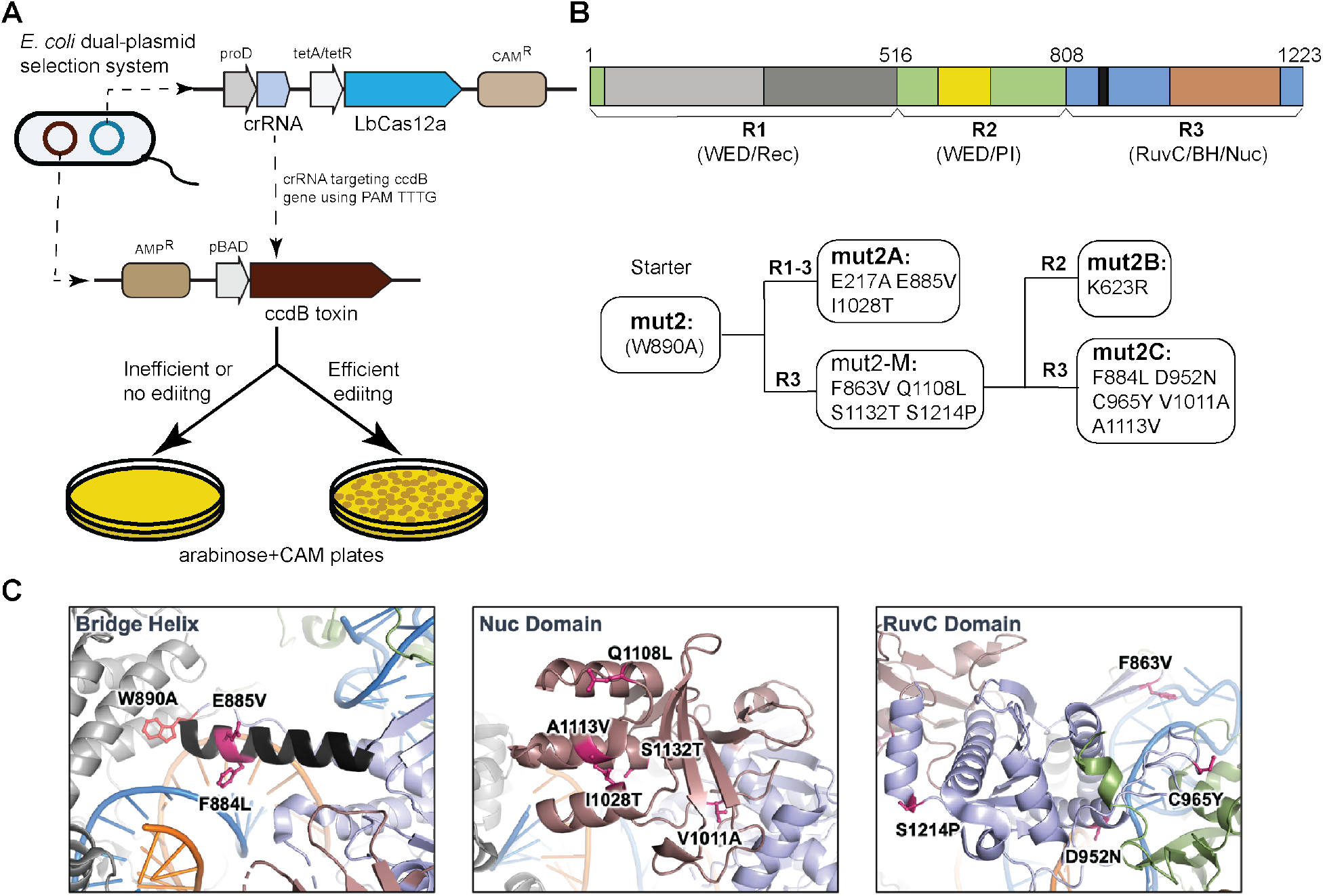
Enhancing *cis*-cleavage activity of mut2 by directed evolution. **A**. Schematic illustration of the positive selection system used for directed evolution of mut2 LbCas12a. **B**. Generation of 3 beneficial mutants from mut2 by directed evolution. Upper panel showing the region (R) divisions of LbCas12a protein used for error-prone polymerase chain reaction (PCR) mutagenesis. Lower panel showing the mutants selected followed each round of mutagenesis. Three beneficial mutants (named mut2A, mut2B, and mut2C) were generated from 2 round of selections. All of the beneficial mutants contain W890A and have the mutations generated in earlier round of selection. **C**. Snapshots of LbCas12a protein structure (PDB ID: 5XUS) highlighting the locations of the beneficial mutations from directed evolution.

The differentiated deleterious effect of W890A in mut2 on *cis*-cleavage and *trans*-cleavage in biochemical analyses made us wonder how this mutation would affect genome editing in mammalian cells. We tested this using neural progenitor cells (NPCs) isolated from Ai9 mice containing a tdTomato transgene under control of a loxP-flanked stop cassette consisting of three repeating transcription terminators (16, 20). LbCas12a-directed removal of the stop cassette would disrupt the terminator and activate expression of tdTomato (Figure 2D). We compared genome editing induced by mut2 versus wild-type LbCas12a using a crRNA that targets the stop cassette and delivering these respective Cas12a ribonucleoproteins (RNPs) into NPCs by nucleofection. Data from fluorescence-activated cell sorting (FACS) of tdTomato-positive cells harvested six days post nucleofection showed that mut2 was 3-10 fold less efficient at inducing genome editing relative to wild-type LbCas12a depending on the nucleofection protocol (Figure 2E). This result is consistent with the 10-fold reduction in the TS cleavage rate observed in our *in vitro* cleavage assays.

### Directed evolution of mutant LbCas12a to improve its genome editing activities

In general, Cas12a has been less widely used in mammalian genome editing when compared to *Streptococcus pyogenes* Cas9 (SpyCas9) (21–23). Improvement of enzymatic activity of LbCas12a could make this group of CRISPR proteins more advantageous for mammalian genome editing. A recent study showed that the activity of Cas12a can be approved through protein engineering (24). In this study, we used directed evolution to select for beneficial mutants derived from mut2. The reduced efficiency of mut2 over wild-type LbCas12 allows for low background that was ideal as a starting point for gain/positive function selection.

We employed a positive bacterial selection scheme, which was initially developed to evolve homing endonucleases (25) and later adapted for CRISPR-based genome editor engineering (26–28). In this approach, positive selection of new Cas12a variants relies on Cas12a-mediated cleavage of a plasmid encoding the toxin-encoding ccdB gene under control of an inducible promoter (Figure 3A). To reduce or minimize the background during selection, we selected a crRNA that targets the ccdB gene at the protospacer containing PAM of TTTG, which is a disfavored PAM *in vitro* by mut2-LbCas12a (Supplementary Figure S3A). When this selection scheme was tested with wild-type LbCas12a, nearly 100% survival of *E. coli* colonies was observed, whereas mut2 produced no bacterial survival (Supplementary Figure S3B). We then conducted directed evolution by randomly mutagenizing the entire mut2-encoding gene at a rate of 6 to 9-nucleotide mutations per kilobase. To simplify the screening process, we divided the entire protein sequence into three regions (R1, R2 and R3) for random mutagenesis carried out with error-prone polymerase chain reaction (PCR) (upper panel, Figure 3B). From two sequential rounds of evolution, three mutants (named mut2A, 2B and 2C) showed nearly 100% bacterial survival under the selection system (Supplementary Figure S3C). Sanger sequencing showed that most of these beneficial mutations in the newly identified mutants are located in the NUC lobe (RuvC, Nuc, and BH domains) (lower panel, Figure 3B, Figure 3C).

We purified these three evolved Cas12a proteins and analyzed their *in vitro* nuclease activities. Kinetic studies of *cis*-cleavage showed that the three new variants displayed improved cutting efficiency, especially on the target strand (Figure 4A,B, Table1, Supplementary Figure S4A, B). Of the three mutants, mut2B and mut2C had *cis*-cleavage activity comparable to the wildtype LbCas12a protein. Specifically, the initial rates of mut2B and mut2C are 6.3 min^−1^ and 7.2 min^−1^ on NTS, and 4.6 min-^1^ and 4.6 min^−1^ on TS, respectively. These values are comparable to those observed for wildtype LbCas12a (7.1 min^−1^ for NTS, 5.7 min^−1^ for TS). We also observed increased *trans*-ssDNase activity for the three mutants (Figure 4C, Table 1, Supplementary Figure S4C), consistent with the inherent partial coupling of *cis* and *trans* activities by the RuvC domain. However, *trans*-ssDNase activity of all three variants still remained 11-100-fold slower relative to the wild-type LbCas12a.

**Figure 4.**
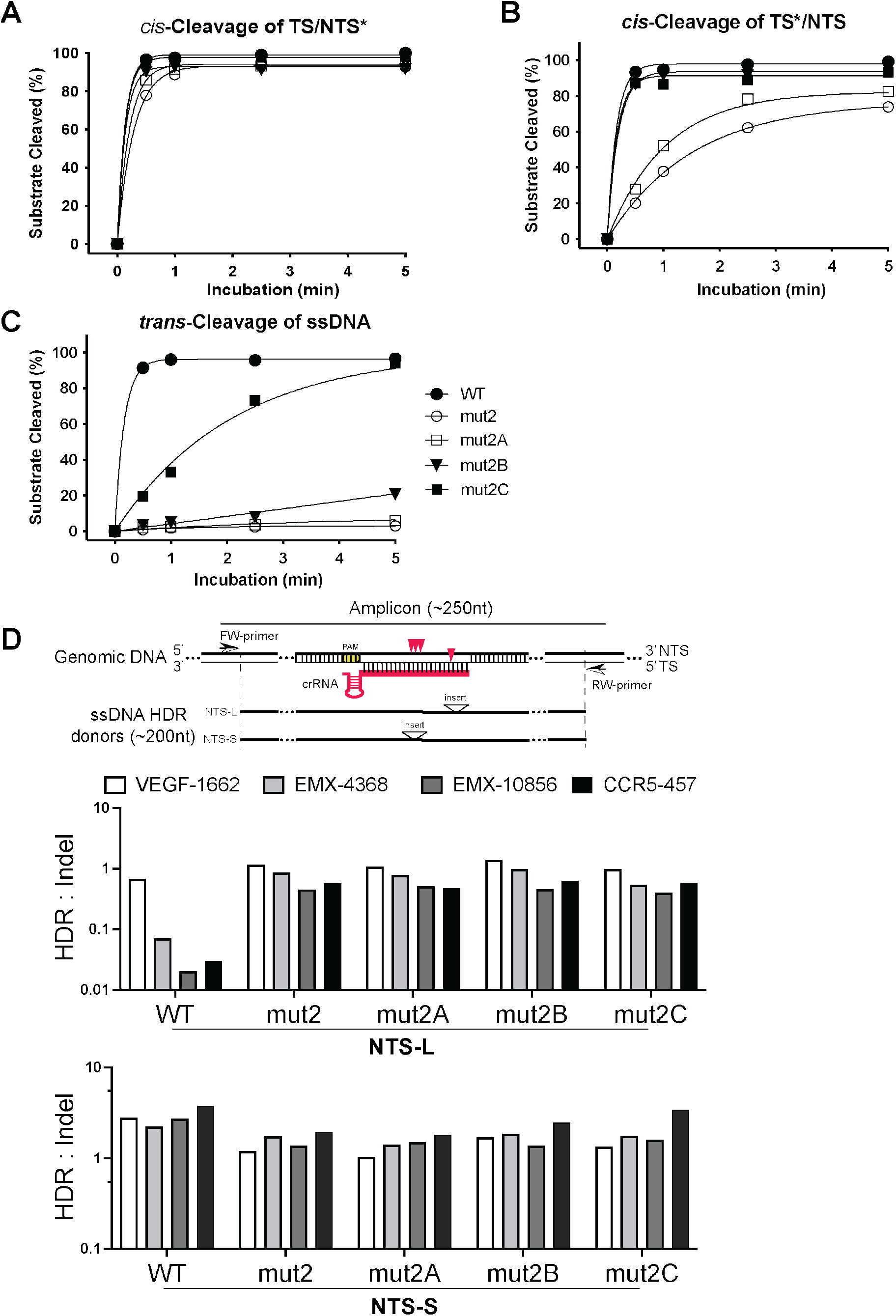
Enhanced activity of mut2A-C *in vitro* and *in cells*. **A-C**. *In vitro* kinetic studies of the cleavage activities on NTS (**A**) and TS (**B**) in a dsDNA as well as the *trans*-cleavage activities (**C**). * indicates labeled strand. All of 3 beneficial mutants from the directed evolution are more active than mut2. mut2B and mut2C display similar *cis*-activities on both strands as wild-type, but their *trans*-activities are much lower. Each data point is averaged from 2-3 parallel experiments. **D**. Genome editing activities of mut2A-C in HEK293 cells. The upper panel shows the nontarget strand (NTS) ssDNA donors of NTS-L and NTS-S defined by the location of inserts from PAM. Specifically, the insert in NTS-L is inserted at 20-24 nt from PAM, while the insert in NTS-S is inserted at 11-14 nt from PAM. Red arrowheads indicate cleavage sites of LbCas12 proteins on target genomic DNA. Insert above the triangle means an exogenous restriction site is inserted as code for calculation of the rate of HDR. The length of ssDNA donors used in this study is less than 200nt and the length of PCR amplicons is less 250 nt. The other two panels show the ratios of HDR:indel calculated from the averaged NGS data from NTS-L (middle panel) and NTS-S (bottom panel). The 3 beneficial mutants from the selection display much better activities in the genome editing in HEK293 cells when donor NTS-L is used. The y-axis is set as log_10_ scale.

**Table 1.** A Summary of rate constant *k* (min^−1^) of all of LbCas12a proteins used in this study

| Protein | WT | mut2 | mut2A | mut2B | mut2C | WT<sup>a</sup> | mut2C-W<sup>a</sup> | mut2C-WF<sup>a</sup> |
| --- | --- | --- | --- | --- | --- | --- | --- | --- |
| *cis*-TS*/NTS | 5.671 | 0.514 | 0.833 | 4.654 | 4.657 | 0.059 | 0.074 | 0.082 |
| *cis*-TS/NTS* | 7.087 | 3.248 | 4.725 | 6.329 | 7.194 | 0.159 | 0.270 | 0.270 |
| *trans*-activity | 5.941 | N.D. | 0.018 | 0.056 | 0.500 | 4.017 | 3.550 | 4.285 |

Note: \*: labeled strand.
a: In these studies, the *cis*-cleavage assays were performed in the reaction containing 20nM protein, 24nM crRNA and 40nM target dsDNA; the *trans*-cleavage assays were carried out in the reaction containing 20nM protein, 24nM crRNA, 30nM dsDNA activator and 120nM ssDNA.
In these studies, fitting curve of a cleavage assay was generated by the formula of $Y = Y_{\max} \times (1 - e^{-kt})$ , where $Y_{\max}$ is the pre-exponential factor, $k$ is the rate constant ( $\text{min}^{-1}$ ), and $t$ is the reaction time (min).

Following biochemical analysis, we also investigated the genome editing activities of these engineered mutants, both for inducing small insertion or deletion mutations (indels) and for homology-directed repair (HDR). We wondered whether Cas12a enzymes with catalytic activity that is robust but biased towards targeted dsDNA cleavage and with minimal *trans*-ssDNase activity might favor genome editing by homology-directed repair (HDR). To test this possibility, we conducted genome-editing experiments using RNPs from the mutants of mut2A, mut2B and mut2C by targeting different genes in HEK293T cells. We created suitable crRNAs for 14 different genomic sites of eight endogenous genes. After 96 hours of cell growth following RNP introduction into cells by nucleofection, genome editing outcomes were preliminarily analyzed and quantified by next-generation sequencing (NGS). Overall, the evolved mutants significantly improved genome editing efficiency when compared to mut2 (Figure 4D and Supplementary Figure S4D1-5). In particular, mut2C displayed similar or better levels of genome editing than the wild-type. To measure rates of HDR, exogenous restriction site (insert)-containing ssDNA oligonucleotides complementary to target or non-target strands were included as repair templates. Interestingly, we observed that these mutants significantly outperform wild-type Cas12a in HDR efficiency with donors containing inserts at PAM-distal region, which results in mismatches with crRNA at 20-24nt away from PAM when Cas12a re-binds to the target site (Figure 4D, Supplementary Figure S4D4,D5).

### Reverse mutations in the bridge helix yield hyper-effective Cas12a genome editors

Biochemical data from this study together with others (15, 19) as well as previous structural analyses (9, 11, 17) suggested that the intact bridge helix (BH) in Cas12a is critical for its activity. The data from this study showing that mut2B and 2C are equal or more active than the wild-type protein biochemically and in cells led us to wonder whether restoring the tryptophan residue (W890) in the BH domain in these mutants might result in even more efficient Cas12a proteins. To test this hypothesis, we selected mut2B and mut2C to study the effect of W890 restoration. As mut2C has another mutation (F884L) in the bridge helix, we also introduced a double reversion, A890W and L884F, into this mutant. The three corresponding variants, mut2B-W, mut2C-W and mut2C-WF, were cloned and purified for functional analysis (Figure 5A).

**Figure 5.**
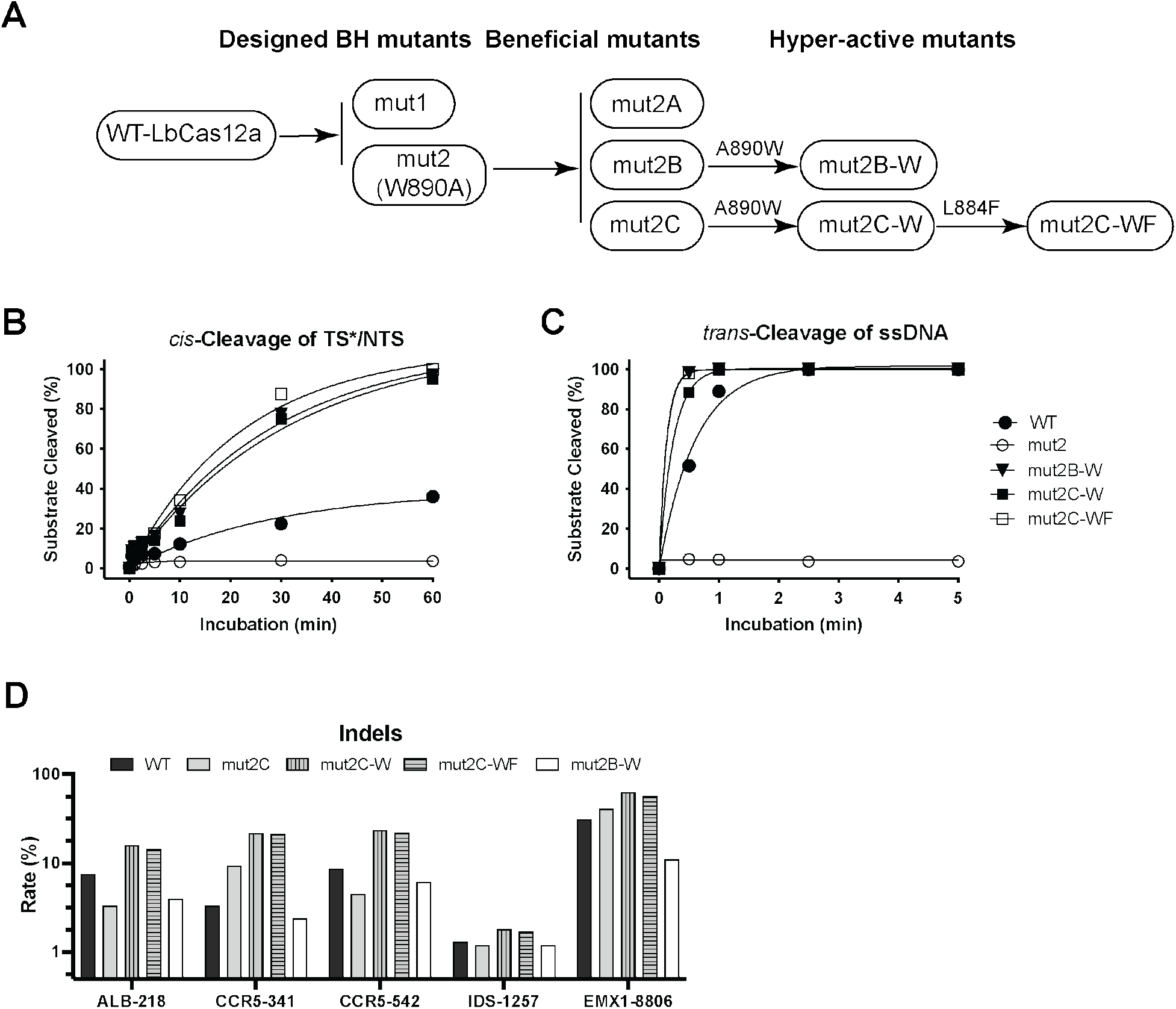
Hyper-effective LbCas12a proteins. **A**. Schematic presentation of the overall pathway for generating beneficial and <u>hyp</u>er-<u>e</u>ffective (HypE) LbCas12a mutants. These 2 HypE-mut2B-W and HypE-mut2C-W mutants are generated by restoring W890 in the corresponding beneficial mutants, respectively. HypE-mut2C-WF is a result of the restorations of both W890 and F884 in mut2C. **B** and **C**. *In vitro* Kinetic studies of the *cis*- and *trans*-activities of the HypE-LbCas12a proteins. Since the HypE-mutants are much more active in both *cis*- and *trans*-activities, we performed the DNA *cis*-cleavage assays in the condition of 20 nM protein, 24 nM crRNA, and 40 nM target-strand-labeled dsDNA. For *trans*-cleavage assays, 120 nM of labeled random ssDNA was used. Each data point is the average of two to three parallel experiments. **D**. Hyper efficient genome editors of HypE-mut2C-W and mut2C-WF. The proteins from HypE-mutants are also hyper efficient in genome editing. The loci selected here are very difficult genomic ones to be edited in HEK293 cells with WT and mutants of mut2A-C. The y-axis is set as log_10_ scale.

We found that the new variants with the restoration of W890 in bridge helix in the evolved mutants are more effective at targeted dsDNA cutting, outperforming the wild-type protein up by 2 fold when comparing initial reaction rates (Figure 5B, Table 1, Supplementary Figure S5A,B). Not surprisingly, their *trans*-cleavage activity against non-specific ssDNA was also significantly increased (Figure 5C, Table1, Supplementary Figure S5C,D). These results suggest that the intact bridge helix together with the mutations identified from directed evolution improve both *cis* and *trans*-cleavage, and the tryptophan of W890 in BH region plays a key role in the sequential activities displayed by Cas12a.

We next tested whether the three improved Cas12a mutants (mut2B-W, mut2C-W and mut2C-WF, Figure 5A) are capable of improved genome editing in mammalian cells. We selected several genomic loci that have proven challenging for genome editing in prior experiments as our editing targets. Analysis of NGS data using these new editors showed that mut2C-W and mut2C-WF produced 2-5-fold improvements in genome editing levels relative to the WT LbCas12a (Figure 5D, Supplementary Figure S5E).

The results from genome editing together with the data from our DNA *in vitro* cleavages (Table 1) clearly showed that these Cas12a mutants of mut2C-W and mut2C-WF are improved Cas12a (iCas12a) editors that can be further developed as robust tools for both genome editing and diagnostics.

## Discussion

In this study, we found that the *cis*-cleavage activity of LbCas12a could be maintained while *trans*-activity was diminished by engineering the bridge helix connecting the REC and NUC lobes of the protein. Key residues in the bridge helix, which are highly conserved among different Cas12a orthologs, interact with the REC-II and RuvC domains and thus contribute to the closed-to-open conformational transitions of the LbCas12a RNP once the target DNA is bound (17, 18). Our study showed that disrupting these interactions by introducing mutations at these sites, such as W890A, E880A, and R883A, affected less on the cleavage of nontarget strand (NTS) but significantly reduced the cleavage of target strand (TS) and even abolished the indiscriminate *trans*-cutting function. The reduction of *cis*-cleavage activity on TS and abolishment of *trans*-cleavage activity displayed in these mutations could be arisen from the reduction of their trimming activity on NTS. It has reported that abolishing or inhibiting NTS trimming activity in Cas12a proteins could eventually slow down the target-strand cleavage and *trans*-cutting events (13, 14). Our results, together with the results from recent studies with FnCas12a (15, 19), highlight the importance of the BH domain in physically bridging and affecting functionally important conformational dynamics of the two major lobes.

Genome-editing experiments from this study showed impaired editing efficiency induced by the mutation of W890A in mut2. The decrease in editing level may stem from two possibilities: (1) protein dynamics are affected by mutation of W890A, resulting in a slower process in target search or trapping of the editor on the cut site which delays the process of recycling; or (2) reduced end-trimming activity on NTS after *cis*-cutting that could revert the cleaved gene to the original state based on DNA repair using base-paired sticky ends. To rescue the gene-editing efficiency in mut2, we carried out a positive selection by directed evolution and obtained three variants from mut2, named as mut2A, mut2B, and mut2C with much more improved nuclease activity. The mutations in these variants are mainly located in the NUC lobe (distributed in RuvC, NUC, and BH domains). The RuvC domain performs the key nuclease function, the BH region regulates protein conformational changes, and the NUC domain is also known to be involved in precisely positioning the target strand of dsDNA prior to its cleavage event (9, 11, 17, 19, 29). Therefore, it is reasonable to speculate that the higher activities displayed in the selected variants might be benefitted from their mutations in NUC lobe which can help tunning the entire NUC lobe to restore the right conformational transitions required for higher *cis*-cleavage activity of Cas12a.

Genome editing data from this study showed that wild-type LbCas12a displayed much lower HDR efficiency when HDR donors containing an exogenous insert at 20-24nt away from PAM were used although its NHEJ efficiency is similar as those showed by the W890A-containing mutants selected from directed evolution. The lower HDR efficiency showed by wild-type LbCas12a could be due to its re-binding and re-cutting of the edited site, whereas such events might be prohibited in these W890A-containing mutants due to their higher sensitivity of mismatches which is supported by the *in vitro* cleavage data from a study with FnCas12a (15). It is highly possible that the higher sensitivity of mismatches displayed in the selected W890A-containing mutants could lead to higher fidelity in genome editing and, therefore, make them to become useful genome editors.

In order to further engineer improved Cas12a gene editors, we reintroduced the tryptophan residue at the end of the bridge helix into the evolved Cas12a variants since the intact bridge helix and helix 1 of the RuvC II domain play structural roles in preserving functionally important conformational states important for target DNA processing (13, 30). Restoration of intact BH domain in the beneficial mutants selected from directed evolution resulted in three hyper effective LbCas12a proteins of mut2B-W, mut2C-W and mut2C-WF. The further improved activity displayed by the these hyper efficient LbCas12a proteins both *in vitro* cleavage on target DNAs and in genome editing in HEK293T cells is very likely arisen from further improvement of their conformational stages. Our results also further supports the importance of the BH domain for the regulation of Cas12a activities and for genome editing efficiency. Overall, the combined engineering and evolutionary strategy described here has created improved Cas12a genome editors and may be applicable to other CRISPR-Cas enzymes in the future.

## Acknowledgments

We thank members of the Doudna lab and the Innovative Genomics Institute for helpful discussions. We would also like to acknowledge Ms. Netravathi Krishnappa (NGS Core Operations Manager and Sequencing Specialist Center for Translational Genomics Innovative Genomics Institute, UC Berkeley) for NGS. This project was funded by grants from the National Science Foundation and by support from the Howard Hughes Medical Institute. Research reported in this publication was supported by the Centers for Excellence in Genomic Science of the National Institutes of Health under award number RM1HG009490. J.A.D. is an investigator of the Howard Hughes Medical Institute.

## Author contributions

Conceptualization, E.M.; K.C.; J.J.L.; B.A.; and J.A.D.; investigation, E.M.; K.C.; H.S.; E.C.S.; K.Z.; J.Y.; supervision, J.A.D.; writing, E.M.; K.C.; H.S.; E.C.S.; and J.A.D.

## REFERENCES

1. Jiang, W. and Marraffini, L.A. (2015) CRISPR-Cas: New Tools for Genetic Manipulations from Bacterial Immunity Systems. Annu Rev Microbiol, 69, 209–228. https://doi.org/10.1146/annurev-micro-091014-104441 http://www.ncbi.nlm.nih.gov/pubmed/26209264

2. Knott, G.J. and Doudna, J.A. (2018) CRISPR-Cas guides the future of genetic engineering. Science, 361, 866–869. https://doi.org/10.1126/science.aat5011 http://www.ncbi.nlm.nih.gov/pmc/articles/PMC6455913

3. Wang, J.Y., Pausch, P. and Doudna, J.A. (2022) Structural biology of CRISPR-Cas immunity and genome editing enzymes. Nat Rev Microbiol, 10.1038/s41579-022-00739-4. https://doi.org/10.1038/s41579-022-00739-4 http://www.ncbi.nlm.nih.gov/pubmed/35562427

4. Zhang, F. and Huang, Z. (2022) Mechanistic insights into the versatile class II CRISPR toolbox. Trends Biochem Sci, 47, 433–450. https://doi.org/10.1016/j.tibs.2021.11.007 http://www.ncbi.nlm.nih.gov/pubmed/34920928

5. Chen, J.S., Ma, E., Harrington, L.B., Da Costa, M., Tian, X., Palefsky, J.M. and Doudna, J.A. (2018) CRISPR-Cas12a target binding unleashes indiscriminate singlestranded DNase activity. Science, 360, 436–439. https://doi.org/10.1126/science.aar6245 http://www.ncbi.nlm.nih.gov/pmc/articles/PMC6628903

6. Gootenberg, J.S., Abudayyeh, O.O., Kellner, M.J., Joung, J., Collins, J.J. and Zhang, F. (2018) Multiplexed and portable nucleic acid detection platform with Cas13, Cas12a, and Csm6. Science, 360, 439–444. https://doi.org/10.1126/science.aaq0179 http://www.ncbi.nlm.nih.gov/pmc/articles/PMC5961727

7. Li, S.-Y., Cheng, Q.-X., Liu, J.-K., Nie, X.-Q., Zhao, G.-P. and Wang, J. (2018) CRISPR-Cas12a has both cis - and trans -cleavage activities on single-stranded DNA. Cell Research, 28, 491–493. https://doi.org/10.1038/s41422-018-0022-x

8. Broughton, J.P., Deng, X., Yu, G., Fasching, C.L., Servellita, V., Singh, J., Miao, X., Streithorst, J.A., Granados, A., Sotomayor-Gonzalez, A., et al. (2020) CRISPR-Cas12-based detection of SARS-CoV-2. Nat Biotechnol, 38, 870–874. https://doi.org/10.1038/s41587-020-0513-4 http://www.ncbi.nlm.nih.gov/pmc/articles/PMC9107629

9. Dong, D., Ren, K., Qiu, X., Zheng, J., Guo, M., Guan, X., Liu, H., Li, N., Zhang, B., Yang, D., et al. (2016) The crystal structure of Cpf1 in complex with CRISPR RNA. Nature, 532, 522–526. https://doi.org/10.1038/nature17944 http://www.ncbi.nlm.nih.gov/pubmed/27096363

10. Yamano, T., Nishimasu, H., Zetsche, B., Hirano, H., Slaymaker, I.M., Li, Y., Fedorova, I., Nakane, T., Makarova, K.S., Koonin, E.V., et al. (2016) Crystal Structure of Cpf1 in Complex with Guide RNA and Target DNA. Cell, 165, 949–962. https://doi.org/10.1016/j.cell.2016.04.003 http://www.ncbi.nlm.nih.gov/pmc/articles/PMC4899970

11. Swarts, D.C., van der Oost, J. and Jinek, M. (2017) Structural Basis for Guide RNA Processing and Seed-Dependent DNA Targeting by CRISPR-Cas12a. Mol Cell, 66, 221–233.e4. https://doi.org/10.1016/j.molcel.2017.03.016 http://www.ncbi.nlm.nih.gov/pmc/articles/PMC6879319

12. Jeon, Y., Choi, Y.H., Jang, Y., Yu, J., Goo, J., Lee, G., Jeong, Y.K., Lee, S.H., Kim, I.-S., Kim, J.-S., et al. (2018) Direct observation of DNA target searching and cleavage by CRISPR-Cas12a. Nat Commun, 9, 2777. https://doi.org/10.1038/s41467-018-05245-x http://www.ncbi.nlm.nih.gov/pmc/articles/PMC6050341

13. Swarts, D.C. and Jinek, M. (2019) Mechanistic Insights into the cis- and trans-Acting DNase Activities of Cas12a. Mol Cell, 73, 589–600.e4. https://doi.org/10.1016/j.molcel.2018.11.021 http://www.ncbi.nlm.nih.gov/pmc/articles/PMC6858279

14. Cofsky, J.C., Karandur, D., Huang, C.J., Witte, I.P., Kuriyan, J. and Doudna, J.A. (2020) CRISPR-Cas12a exploits R-loop asymmetry to form double-strand breaks. Elife, 9, e55143. https://doi.org/10.7554/eLife.55143 http://www.ncbi.nlm.nih.gov/pmc/articles/PMC7286691

15. Wörle, E., Jakob, L., Schmidbauer, A., Zinner, G. and Grohmann, D. (2021) Decoupling the bridge helix of Cas12a results in a reduced trimming activity, increased mismatch sensitivity and impaired conformational transitions. Nucleic Acids Res, 49, 5278–5293. https://doi.org/10.1093/nar/gkab286 http://www.ncbi.nlm.nih.gov/pmc/articles/PMC8136826

16. Staahl, B.T., Benekareddy, M., Coulon-Bainier, C., Banfal, A.A., Floor, S.N., Sabo, J.K., Urnes, C., Acevedo Munares, G., Ghosh, A. and Doudna, J.A. (2017) Efficient genome editing in the mouse brain by local delivery of engineered Cas9 ribonucleoprotein complexes. Nat Biotechnol, 35, 431–434. https://doi.org/10.1038/nbt.3806 http://www.ncbi.nlm.nih.gov/pmc/articles/PMC6649674

17. Stella, S., Mesa, P., Thomsen, J., Paul, B., Alcón, P., Jensen, S.B., Saligram, B., Moses, M.E., Hatzakis, N.S. and Montoya, G. (2018) Conformational Activation Promotes CRISPR-Cas12a Catalysis and Resetting of the Endonuclease Activity. Cell, 175, 1856–1871.e21. https://doi.org/10.1016/j.cell.2018.10.045 http://www.ncbi.nlm.nih.gov/pubmed/30503205

18. Swarts, D.C. (2019) Making the cut(s): how Cas12a cleaves target and non-target DNA. Biochemical Society Transactions, 47, 1499–1510. https://doi.org/10.1042/BST20190564

19. Parameshwaran, H.P., Babu, K., Tran, C., Guan, K., Allen, A., Kathiresan, V., Qin, P.Z. and Rajan, R. (2021) The bridge helix of Cas12a imparts selectivity in cis-DNA cleavage and regulates trans-DNA cleavage. FEBS Lett, 595, 892–912. https://doi.org/10.1002/1873-3468.14051 http://www.ncbi.nlm.nih.gov/pmc/articles/PMC8044059

20. Madisen, L., Zwingman, T.A., Sunkin, S.M., Oh, S.W., Zariwala, H.A., Gu, H., Ng, L.L., Palmiter, R.D., Hawrylycz, M.J., Jones, A.R., et al. (2010) A robust and high-throughput Cre reporting and characterization system for the whole mouse brain. Nat Neurosci, 13, 133–140. https://doi.org/10.1038/nn.2467 http://www.ncbi.nlm.nih.gov/pmc/articles/PMC2840225

21. Zetsche, B., Heidenreich, M., Mohanraju, P., Fedorova, I., Kneppers, J., DeGennaro, E.M., Winblad, N., Choudhury, S.R., Abudayyeh, O.O., Gootenberg, J.S., et al. (2017) Multiplex gene editing by CRISPR-Cpf1 using a single crRNA array. Nat Biotechnol, 35, 31–34. https://doi.org/10.1038/nbt.3737 http://www.ncbi.nlm.nih.gov/pmc/articles/PMC5225075

22. Chow, R.D., Wang, G., Ye, L., Codina, A., Kim, H.R., Shen, L., Dong, M.B., Errami, Y. and Chen, S. (2019) In vivo profiling of metastatic double knockouts through CRISPR-Cpf1 screens. Nat Methods, 16, 405–408. https://doi.org/10.1038/s41592-019-0371-5 http://www.ncbi.nlm.nih.gov/pmc/articles/PMC6592845

23. Duarte, F. and Déglon, N. (2020) Genome Editing for CNS Disorders. Front Neurosci, 14, 579062. https://doi.org/10.3389/fnins.2020.579062 http://www.ncbi.nlm.nih.gov/pmc/articles/PMC7642486

24. Guo, L.Y., Bian, J., Davis, A.E., Liu, P., Kempton, H.R., Zhang, X., Chemparathy, A., Gu, B., Lin, X., Rane, D.A., et al. (2022) Multiplexed genome regulation in vivo with hyper-efficient Cas12a. Nat Cell Biol, 24, 590–600. https://doi.org/10.1038/s41556-022-00870-7 http://www.ncbi.nlm.nih.gov/pmc/articles/PMC9035114

25. Chen, Z. and Zhao, H. (2005) A highly sensitive selection method for directed evolution of homing endonucleases. Nucleic Acids Res, 33, e154. https://doi.org/10.1093/nar/gni148 http://www.ncbi.nlm.nih.gov/pmc/articles/PMC1253837

26. Hu, J.H., Miller, S.M., Geurts, M.H., Tang, W., Chen, L., Sun, N., Zeina, C.M., Gao, X., Rees, H.A., Lin, Z., et al. (2018) Evolved Cas9 variants with broad PAM compatibility and high DNA specificity. Nature, 556, 57–63. https://doi.org/10.1038/nature26155 http://www.ncbi.nlm.nih.gov/pmc/articles/PMC5951633

27. Lee, J.K., Jeong, E., Lee, J., Jung, M., Shin, E., Kim, Y., Lee, K., Jung, I., Kim, D., Kim, S., et al. (2018) Directed evolution of CRISPR-Cas9 to increase its specificity. Nat Commun, 9, 3048. https://doi.org/10.1038/s41467-018-05477-x

28. Kleinstiver, B.P., Sousa, A.A., Walton, R.T., Tak, Y.E., Hsu, J.Y., Clement, K., Welch, M.M., Horng, J.E., Malagon-Lopez, J., Scarfò, I., et al. (2019) Engineered CRISPR–Cas12a variants with increased activities and improved targeting ranges for gene, epigenetic and base editing. Nature Biotechnology, 37, 276–282. https://doi.org/10.1038/s41587-018-0011-0

29. Stella, S., Alcón, P. and Montoya, G. (2017) Structure of the Cpf1 endonuclease R-loop complex after target DNA cleavage. Nature, 546, 559–563. https://doi.org/10.1038/nature22398 http://www.ncbi.nlm.nih.gov/pubmed/28562584

30. Wörle, E., Newman, A., Burgio, G. and Grohmann, D. (2022) Allosteric activation of CRISPR-Cas12a requires the concerted movement of the bridge helix and helix 1 of the RuvC II domain Biochemistry. https://doi.org/10.1101/2022.03.15.484427

