## Supplementary Figures for "Improved Genome Editing by an Engineered CRISPR-Cas12a"

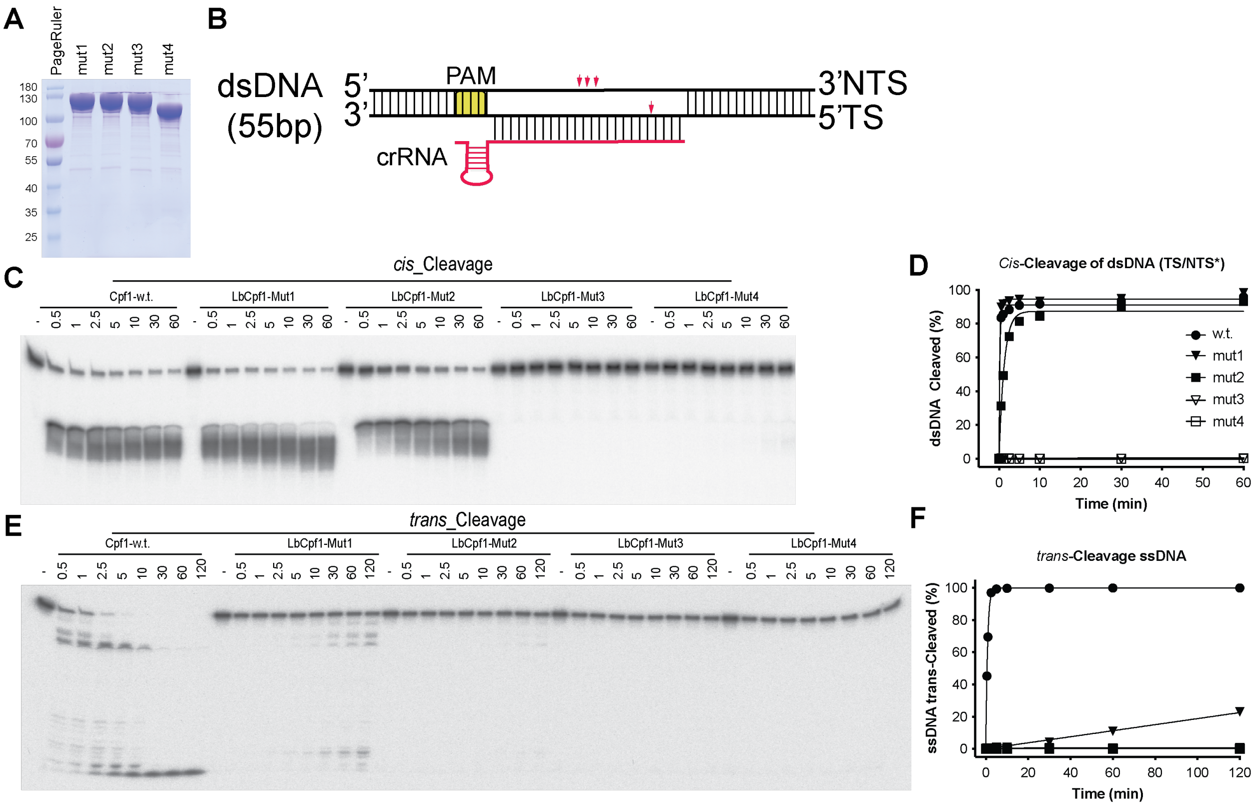


**Supplementary Figure 1**. Importance of the bridge helix (BH) of LbCas12a protein in regulating its nuclease activities. **A**. Purification of 4 LbCas12a mutant proteins. **B**. Schematic presentation of a target dsDNA (55bp) and 43nt CRISPR RNA (crRNA) containing a 19-base direct repeat region (loop domain) and a 24-base protospacer region complementary to the center of a 55-base pair (bp) dsDNA substrate in which the target region flanks the protospacer adjacent motif (PAM) of TTTA. **C**,**D**. *In vitro* kinetic studies of *cis*-cleavage activities by wild-type (WT) and four designed mutants, mut1-4. Each data point is the average of two to three parallel experiments. C shows actual gel image and D is quantification from C. **E**,**F**. *In vitro* kinetic studies of *trans*-cleavage activities by wild-type (WT) and four designed mutants, mut1-4. Each data point is the average of two to three parallel experiments. E shows actual gel image and F is quantification from E.

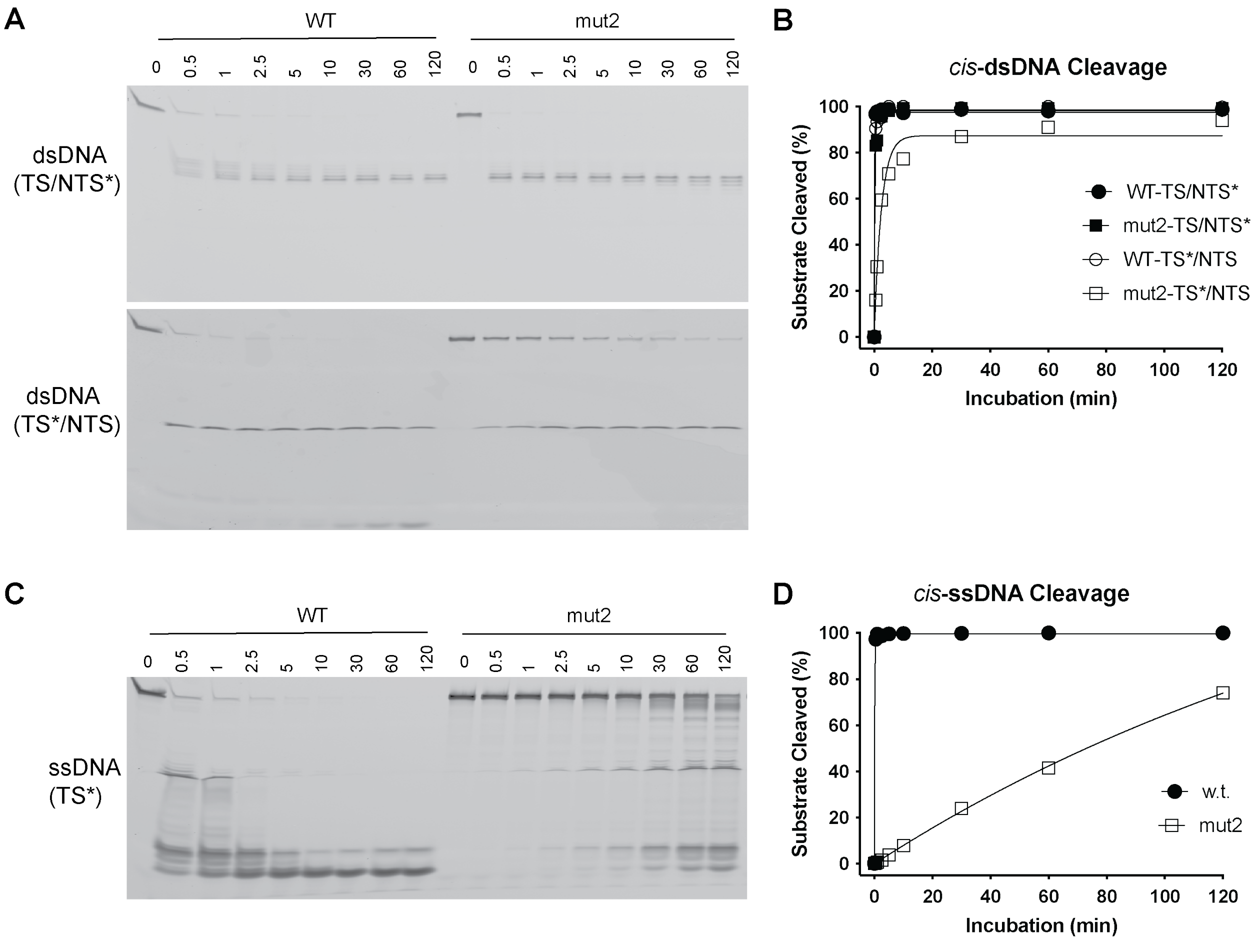


**Supplementary Figure 2**. Kinetic studies of mutational effect of W890A on the nuclease activities of LbCas12a proteins. **A**,**B**. *cis*-cleavage assays of dsDNA. A shows 2 actual images of the *cis*-cleavage activities on dsDNA. In NTS cleavage, trimming activity can be clearly seen with wild-type protein, but no with mut2 protein. B is quantifications from A. **C**,**D**. *cis*-cleavage assays of ssDNA. Actual image of the *cis*-cleavage activities on ssDNA (C) and their quantifications (D). TS=target strand; NTS= nontarget strand; *=labelled strand. mut2 is W890A point mutation of LbCas12a.

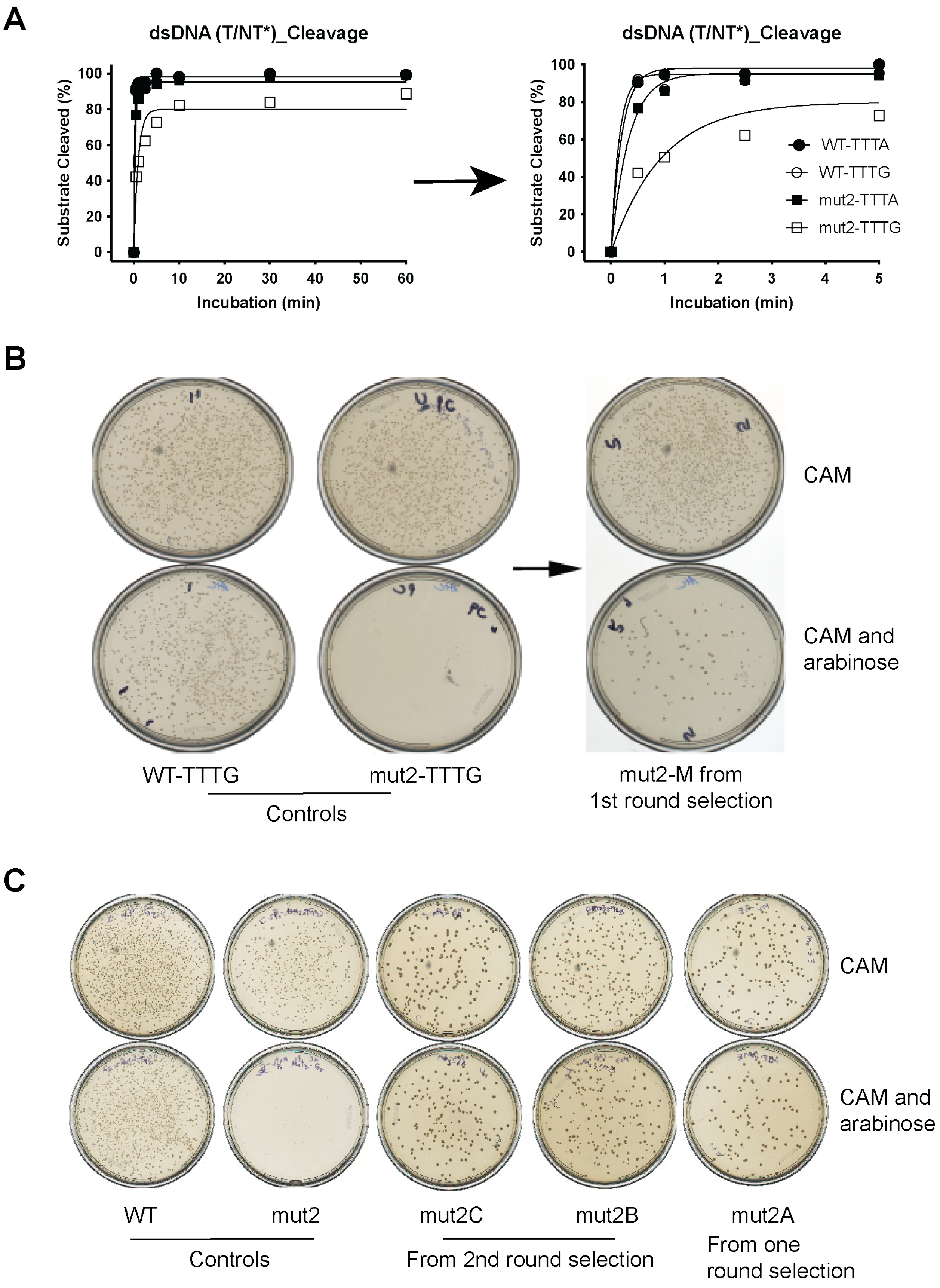

A
dsDNA (T/NT*)_Cleavage
Substrate Cleaved (%)
Incubation (min)
dsDNA (T/NT*)_Cleavage
Substrate Cleaved (%)
Incubation (min)
WT-TTTA
WT-TTTG
mut2-TTTA
mut2-TTTG
B
CAM
CAM and arabinose
WT-TTTG
mut2-TTTG
Controls
mut2-M from 1st round selection
C
CAM
CAM and arabinose
WT
mut2
Controls
mut2C
mut2B
From 2nd round selection
mut2A
From one round selection

**Supplementary Figure 3**. More active variants of mut2 selected by directed evolution. **A**. Effects of mut2 (W890A) on PAM usage. mut2 disfavors TTTG PAM which is used for the directed evolution selection. **B**. First round selection of directed evolution. Selected a candidate, named mut2M, with mutations of F863V, W890A, Q1108L, S1132T and S1214P. **C**. Second round selection using mut2M. Two variants with higher activity were selected: mut2B: K623R, F863V, W890A, Q1108L, S1132T, S1214P and mut2C: F863V, F884L, W890A, D952N, C965Y, V1011A, Q1108L, A1113V, S1132T, S1214P. mut2A was selected in one round error-prone PCR of whole LbCas12a: E217A, E885V, W890A, I1028T;

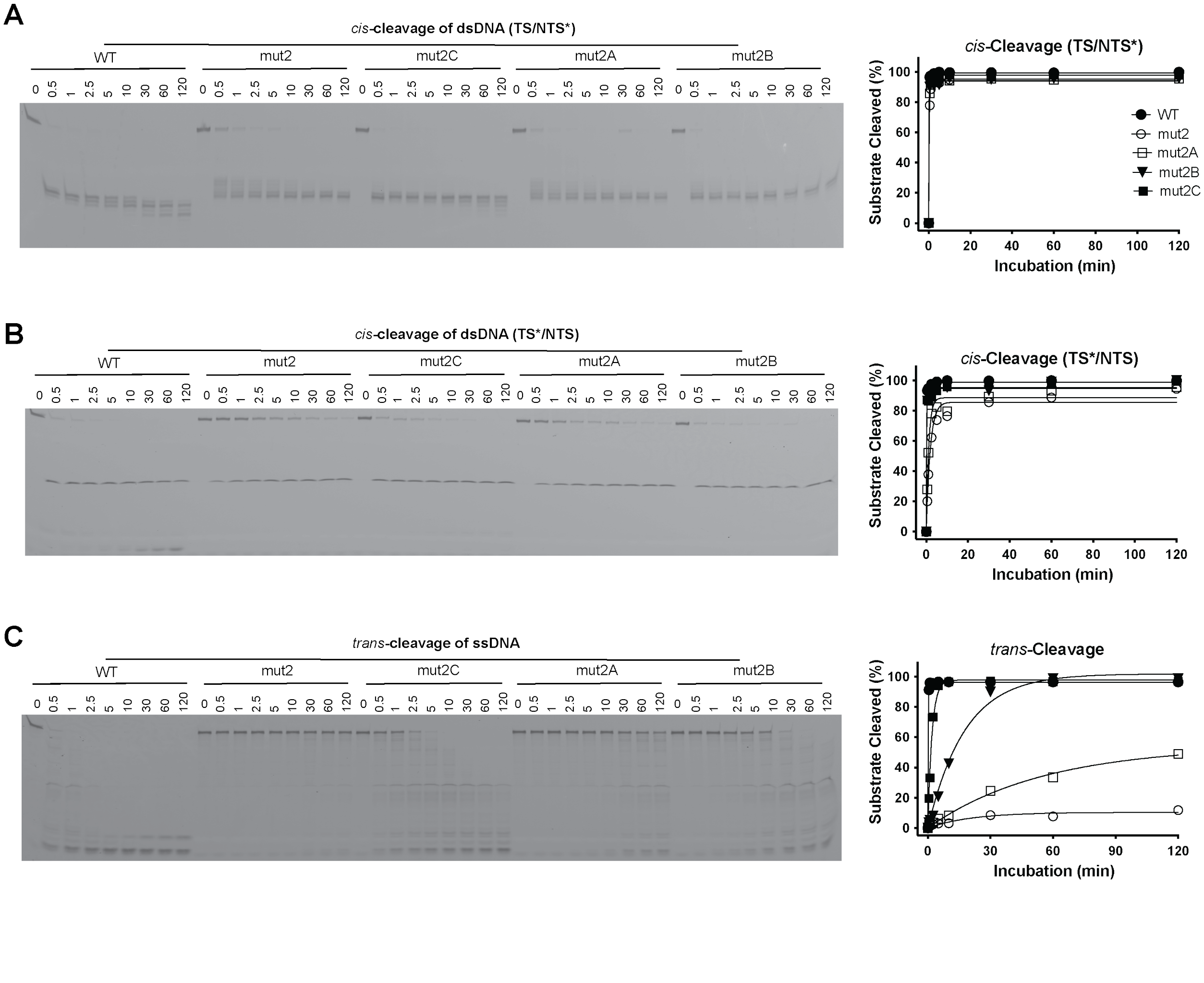

D1

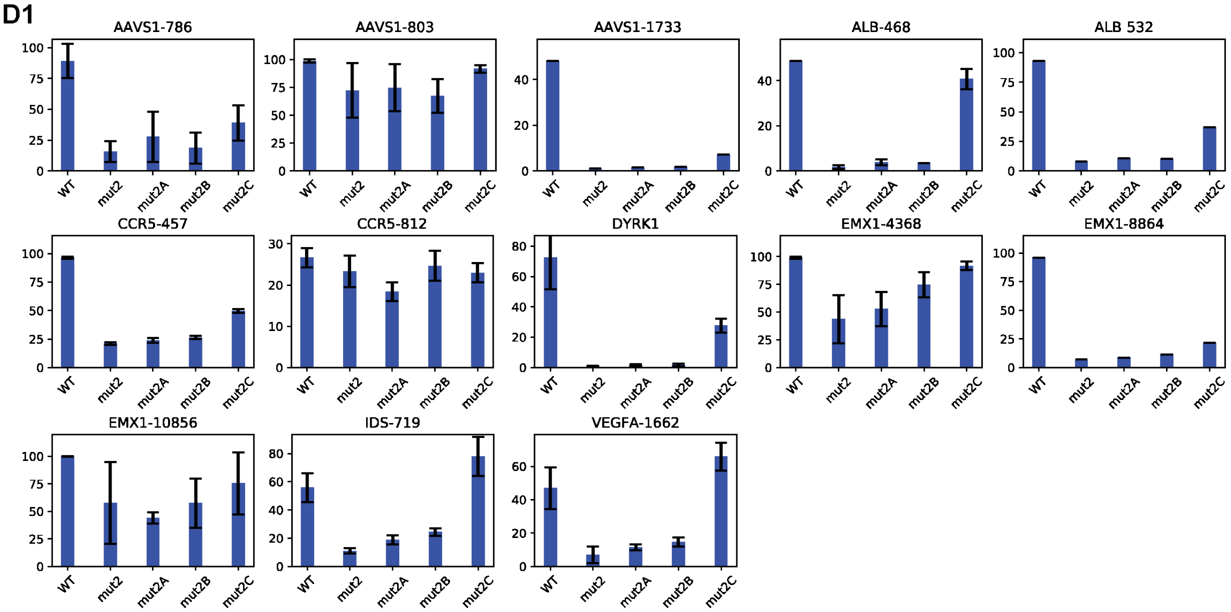


D2

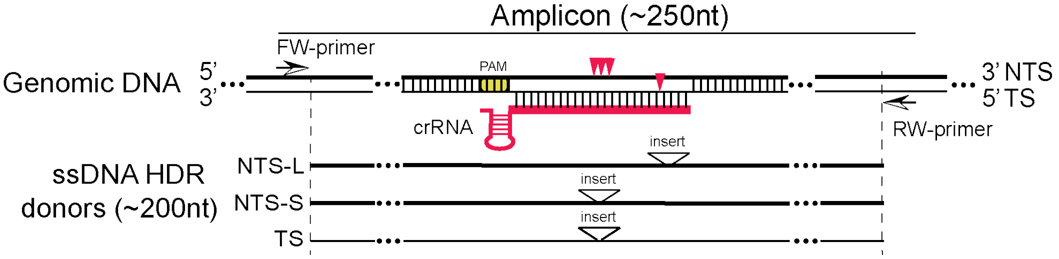


D3

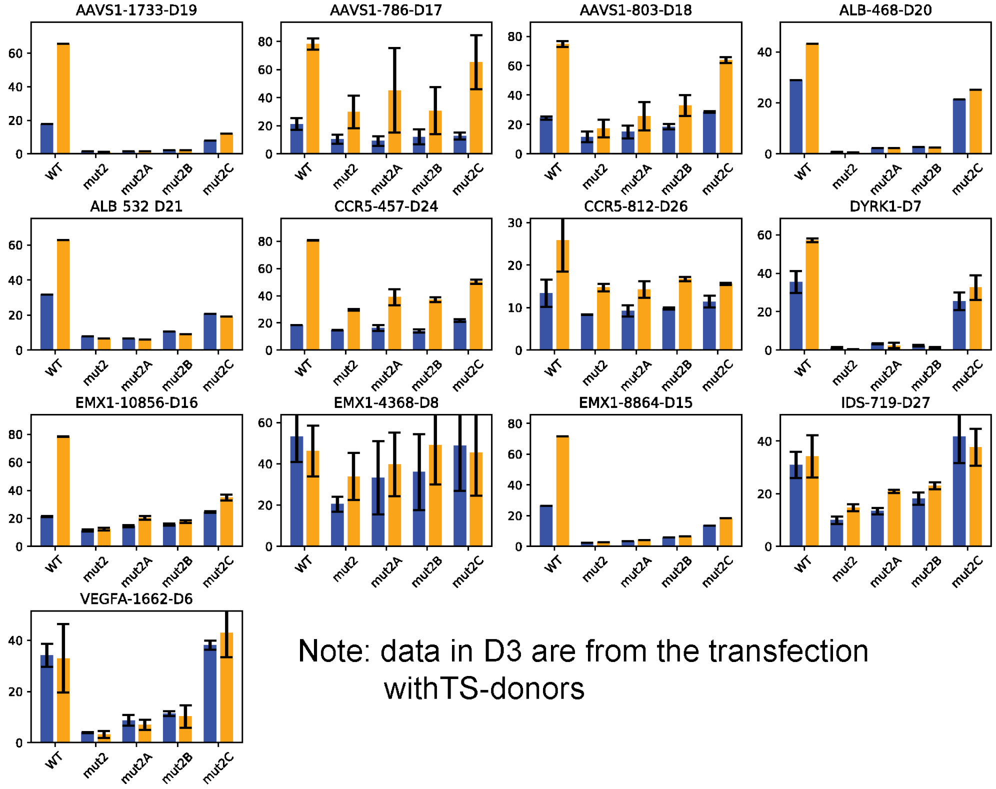


Note: data in D3 are from the transfection withTS-donors

**D4**

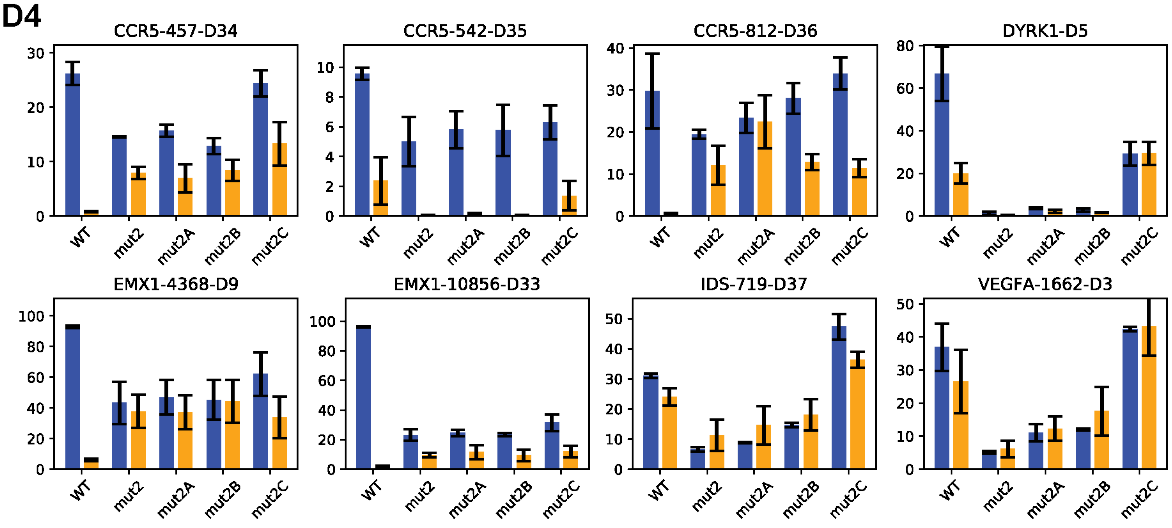


**D5**

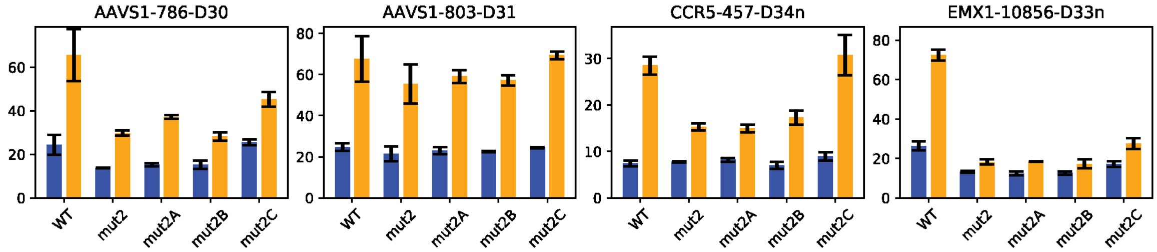


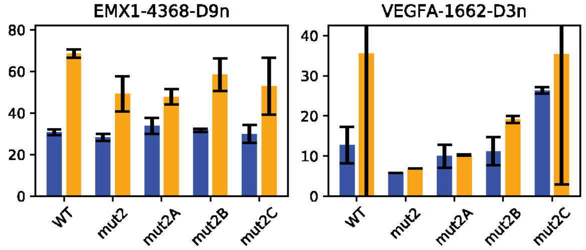


Note: data in D4 are from the transfection with NTS-L donors; while data in D5, NTS-S donors.

**Supplementary Figure 4**. Enhanced activity of mut2A-C *in vitro*. **A-C**. *In vitro* kinetic studies of the cleavage activities on NTS (**A**) and TS (**B**) in a dsDNA as well as the *trans*-cleavage activities (**C**). * indicates labeled strand. Left panels are actual cleavage images and left panels are the corresponding quantifications. All of 3 beneficial mutants from the directed evolution are more active than mut2. mut2B and mut2C display similar *cis*-activities on both strands as wild-type, but their *trans*-activities are still much lower. Each data point is averaged from 2-3 parallel experiments. **D**1-5. Genome editing in HEK293T cells by the new variants selected from mut2 by directed evolution. **D**1 shows the rate (%) of NHEJ when the cells are transfected with only LbCas12a RNPs; **D**2 is schematic presentation showing the ssDNA donors for TS and NTS. For NTS donors, two kinds of NTS donors were designed: NTS-L and NTS-S defined by the location of inserts from PAM. Specifically, the insert in NTS-L is located at 20-24nt from PAM, while the insert in NTS-S is located at 11-14nt from PAM. Red arrowheads indicate cleavage sites of LbCas12 proteins on target genomic DNA. Insert above the triangle means an exogenous restriction site is inserted as code for calculation of the rate of HDR. The length of ssDNA donors used in this study is less than 200nt and the length of PCR amplicons is less 250nt; **D**3 shows the rates (%) of NHJEs and HDRs when the cells are transfected with both LbCas12a RNPs and TS ssDNA donors; **D**4 shows the rates (%) of NHJEs and HDRs when the cells are transfected with both LbCas12a RNPs and NTS-L ssDNA donors; **D**5 shows the rates (%) of NHJEs and HDRs when the cells are transfected with both LbCas12a RNPs and NTS-S ssDNA donors.

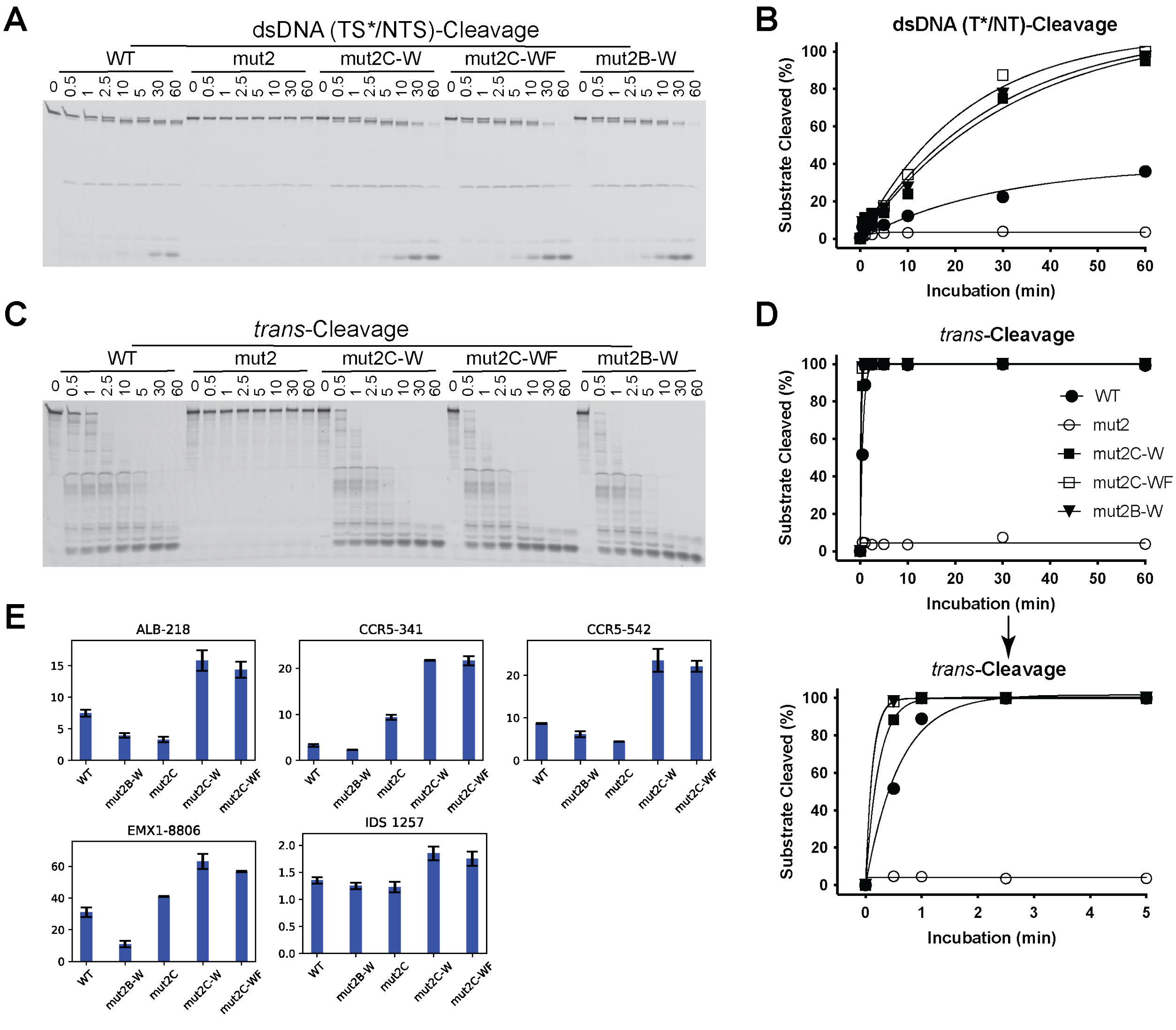

**Supplementary Figure 5**. Hyper-effective LbCas12a proteins. Restoration of tryptophan at W890 in the 2 beneficial mutants (mut2B and mut2C) generates hyper effective (HypE) mutants: mut2B-W, mut2C-W and mut2C-WF which contains two conversions F884 and W890. **A**,**B**. *In vitro* Kinetic studies of the *cis*-activities of the HypE-LbCas12a proteins. A is an actual clevage image; B is quantification of A. The HypE-mutants are much more active in *cis*-dsDNA cleavage. Here, the *cis*-cleavage assays were carried out with 20nM protein, 24nM crRNA, and 40nM target-strand-labeled dsDNA. Each data point is the average of two to three parallel experiments. **C**,**D**. *In vitro* Kinetic studies of the *trans*-cleavage activities of the HypE-LbCas12a proteins. C is an actual cleavage image; D is quantification of C. Here, the *cis*-cleavage assays were carried out with 20nM protein, 24nM crRNA, 40nM activator dsDNA and 120nM of *trans*-ssDNA. **E**. Genome editing in HEK293T cells by hyper efficient LbCas12a proteins. Rate (%) of NHEJs are presented. These loci are very challenging to be edited by wild-type LbCas12a.
